# Harnessing Expressed Single Nucleotide Variation and Single Cell RNA Sequencing to Define Immune Cell Chimerism in the Rejecting Kidney Transplant

**DOI:** 10.1101/2020.03.10.986075

**Authors:** Andrew F. Malone, Haojia Wu, Catrina Fronick, Robert Fulton, Joseph P. Gaut, Benjamin D. Humphreys

## Abstract

In solid organ transplantation, donor derived immune cells are assumed to decline with time after surgery. Whether donor leukocytes persist within kidney transplants or play any role in rejection is unknown, however, in part because of limited techniques for distinguishing recipient and donor cells. To address this question, we performed paired whole exome sequencing of donor and recipient DNA and single cell RNA sequencing (scRNA-seq) of 5 human kidney transplant biopsy cores. Exome sequences were used to define single nucleotide variations (SNV) across all samples. By analyzing expressed SNVs in the scRNA-seq dataset we could define recipient vs. donor cell origin for all 81,139 cells. The leukocyte donor to recipient ratio varied with rejection status for macrophages and with time post-transplant for lymphocytes. Recipient macrophages were characterized by inflammatory activation and donor macrophages by antigen presentation and complement signaling. Recipient origin T cells expressed cytotoxic and pro-inflammatory genes consistent with an effector cell phenotype whereas donor origin T cells are likely quiescent expressing oxidative phosphorylation genes relative to recipient T cells. Finally, both donor and recipient T cell clones were present within the rejecting kidney, suggesting lymphoid aggregation. Our results indicate that donor origin macrophages and T cells have distinct transcriptional profiles compared to their recipient counterparts and donor macrophages can persist for years post transplantation. This study demonstrates the power of this approach to accurately define leukocyte chimerism in a complex tissue such as the kidney transplant coupled with the ability to examine transcriptional profiles at single cell resolution.

## Introduction

Single cell RNA sequencing (scRNA-seq) technologies offer the ability to measure the transcriptomes of many thousands of cells simultaneously. The resulting datasets have revealed unexpected heterogeneity between cell types and states and new biologic insights.(1, 2) Natural genetic variations within the coding region of the genome is also captured by scRNA-seq. Using germ line coding sequence data as a reference this information can be harnessed to identify the origin of each cell from a sample of cells of mixed origin.(3)

Kidney transplantation offers the greatest survival benefit for patients with end stage kidney disease (ESKD). It was appreciated very early on that acquired tolerance of donor tissues was associated with leukocyte chimerism.(4) In the 1960s long term immunosuppression free graft survival was not uncommon in living related transplants.(5) However, with the advent of modern immunosuppression short-term outcomes improved to above 90% at one year. Thus, interest in leukocyte chimerism waned and immunosuppression free transplantation became a rare occurrence. A renewed interest in the relevance of donor leukocyte chimerism has occurred since the 1990s.(6) The existence of solid organ transplant associated graft vs. host disease (GvHD) has suggested that donor derived immune cells can persist and cause pathology years later but this is extremely rare in kidney, with only six cases reported.(7–9) Furthermore, donor derived alveolar macrophages can survive in lung transplants for over 3 years.(10) Recently, investigators used HLA-disparate lung transplant recipients and FACS sorting to show that donor derived T cells persist in lung transplantation.(11) Also, adoptive transfer of donor origin T cells has been shown to modulate the alloimmune response.(12, 13) Other studies report that tissue resident macrophages are replaced by monocyte-derived cells post injury.(14, 15) Whether tissue resident leukocytes such as macrophages persist and what role they play in kidney is completely unknown. We hypothesized that the immune cell infiltrate in the human kidney transplant is composed of donor and recipient origin cells and that the transcriptional patterns of donor origin immune cells differ from those of recipient cells.

Here we describe scRNA-seq analysis of 5 kidney transplant biopsies from patients with acute ABMR or non-rejection acute kidney injury (AKI). We identified all the major kidney cell types and two main immune cell types, lymphocytes and macrophages. The lymphocyte cluster contained T cells, B cells and plasma cells. Using whole exome sequence from recipient and donor for each biopsy sample, we could establish the recipient or donor origin of all of these cells based on expressed single nucleotide polymorphisms. Donor origin cells made up a substantial proportion of macrophages and lymphocytes. The proportion of donor to recipient cells varies with rejection status for macrophages and with time post-transplant for lymphocytes. Donor origin macrophages express antigen presentation and complement pathway genes. Donor origin T cell gene expression correlates with the non-rejecting state. Oxidative phosphorylation genes are differentially expressed in donor origin T cells compared to recipient T cells suggesting donor T cells are in a quiescent state.(16) This study demonstrates the power of this approach applied to the study of leukocyte chimerism in organ transplantation. Further analysis of donor and host chimerism by scRNA-seq should greatly advance our understanding of the roles donor and recipient immune cells play in the alloimmune response.

## Results

Kidney allograft tissue was taken by needle core biopsy from 3 patients with ABMR and 2 patients with a non-rejection cause of AKI. All patients diagnosed with ABMR had donor specific antibodies (DSA) at the time of biopsy. One patient had received an ABO incompatible kidney transplant and anti-A isoantibody (DSA) was elevated at the time of biopsy. The other two patients with ABMR had at least one anti-HLA antibody (DSA) with mean fluorescence intensities (mfi) of at least 5000. In one patient with a non-rejection biopsy diagnosis a low mfi (< 2000) anti-HLA antibody was present at the time of diagnosis. Biopsies were performed at 5 days up to 7 years post transplantation. All patients received anti-thymocyte globulin and methylprednisolone at the time of transplant followed by maintenance immunosuppression of tacrolimus, mycophenolic acid and prednisone. One rejecting patient and one non-rejecting patient received additional anti-CD20 antibody infusion at the time of transplantation (Table S1).

### scRNA-seq of allograft biopsies resolves the major kidney and immune cell types

A total of 81,139 cells passed filters with an average of 1124 genes and 2497 transcripts per cell. Unsupervised clustering analysis identified eight individual clusters of kidney cells and two immune cell clusters. One cluster was identified as stressed proximal tubular cells as they differentially expressed genes regulating apoptosis and oxidative stress (Fig. S1). We identified 5027 macrophages and 3647 lymphocytes in the same clustering analysis (Fig. 1). There was minimal batch effect in this dataset (Fig. S2).

**Figure 1.**
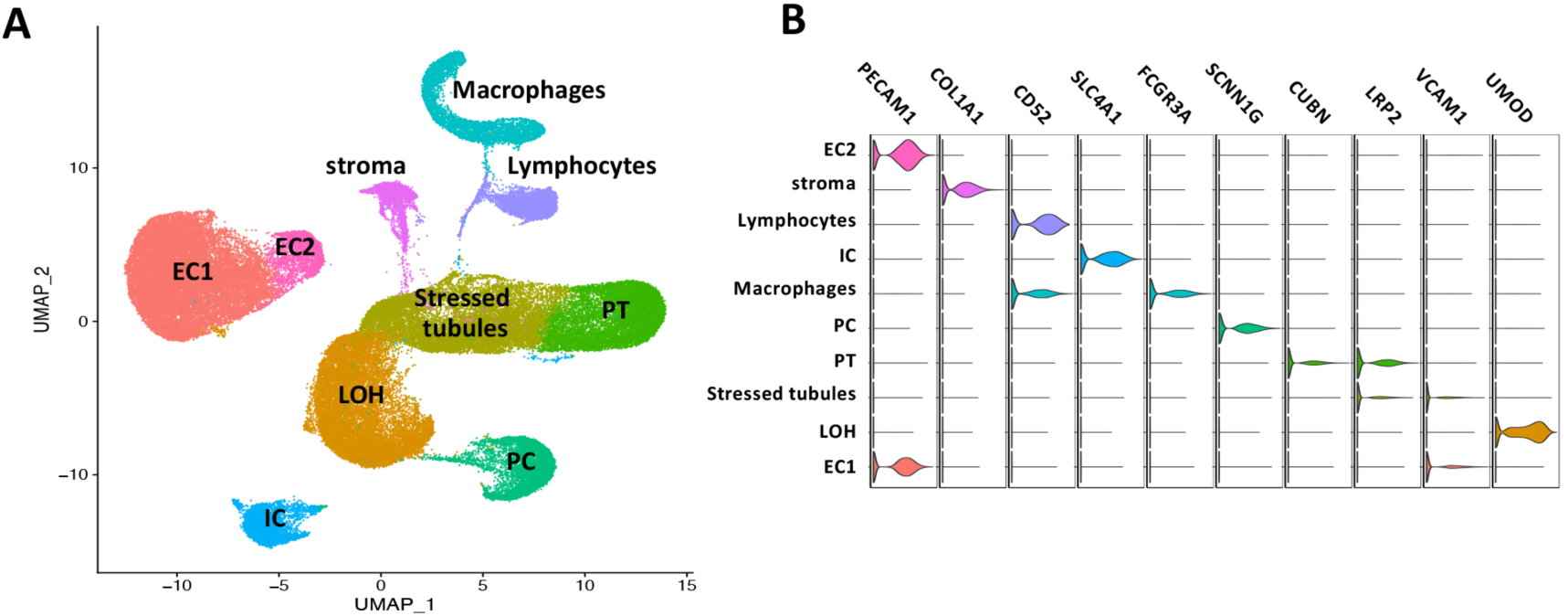
Visualization of 81,139 Cells from 5 Kidney Allograft Biopsies Representing all Major Kidney and Immune Cell Types. A) UMAP plot of 81,139 cells from 5 kidney allograft biopsies. All the main kidney cell types are represented including a stressed tubular cells, loop of Henle (LOH), proximal tubule (PT), principal cells (PC), intercalated cells (IC), endothelial cells (EC), stromal cells, macrophages and lymphocytes. B) Violin plot demonstrating marker genes for major cell types represented in the whole dataset and UMAP plot.

### Donor and recipient immune cell chimerism in the kidney allograft

Whole-exome sequencing (WES) was performed on the 10 individuals from the five donor recipient pairs for each biopsy. A single variant call format (.vcf) file containing an average of 120,896 coding SNVs with high population minor allele frequency (maf > 5%) was constructed from WES data for each donor and recipient pair. The demuxlet pipeline(3) was applied to each of the 5 biopsies in this dataset using the .vcf file as a reference to determine donor or recipient origin of each cell (Fig. 2A). Despite manual filtering and removal of doublet cells from the single cell RNA-seq data (see supplementary methods) from each biopsy the demuxlet tool identified 3 - 10.5% of cells as doublets which allowed for improved quality control of the dataset. The macrophage cluster was comprised of 3798 recipient cells and 1229 donor origin cells and the lymphocyte cluster comprised of 1528 recipient cells and 2119 donor cells (Fig. 2B).

**Figure 2.**
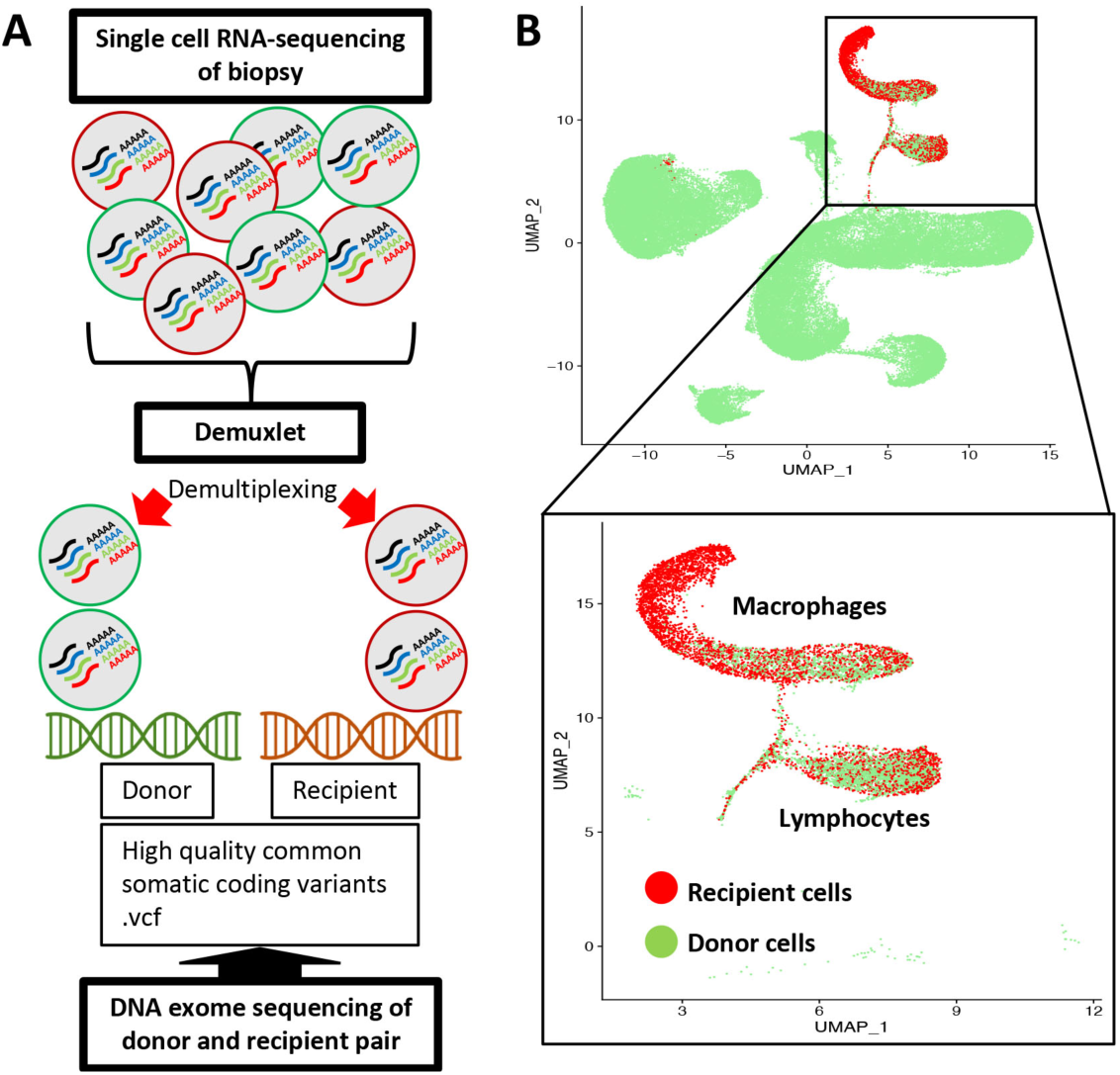
There is Donor and Recipient Chimerism of Macrophages and Lymphocytes in the Transplanted Kidney. A) The Demuxlet pipeline used to resolve the origin of cells from each biopsy. Kang HM et al. *Nat Biotechnol.* 2018;36(1):89-94. B) UMAP visualization of 81,139 cells grouped by recipient (red) or donor (green) origin. Inset highlights macrophages (5027 cells) and lymphocytes (3647 cells) grouped by recipient or donor origin.

The ratio of donor to recipient origin macrophages varied according to rejection status with approximately equal proportions of donor and recipient cells in the two non-rejecting samples and almost complete dominance of recipient cells in rejecting biopsies (Fig. 3A). The absolute number of donor and recipient origin macrophages in rejecting samples ranged between 4 – 39 cells and 514 – 1258 cells, respectively. In non-rejecting samples donor and recipient origin macrophage numbers ranged between 217 – 964 and 389 – 674, respectively. This difference occurred independent of time post-transplant. As an example, there were a significant proportion (35.8%) of donor macrophages in the non-rejecting allograft at 28 days whereas in the rejecting biopsy at day 11 post-transplant recipient macrophages dominate (97%) (Fig. 3B). In an independent dataset donor origin macrophage numbers recovered post rejection treatment increasing from 3% of total macrophages to 11.7% one month post treatment (Fig. S3). Furthermore, absolute numbers of donor macrophages increased post treatment from 96 to 100 cells.

**Figure 3.**
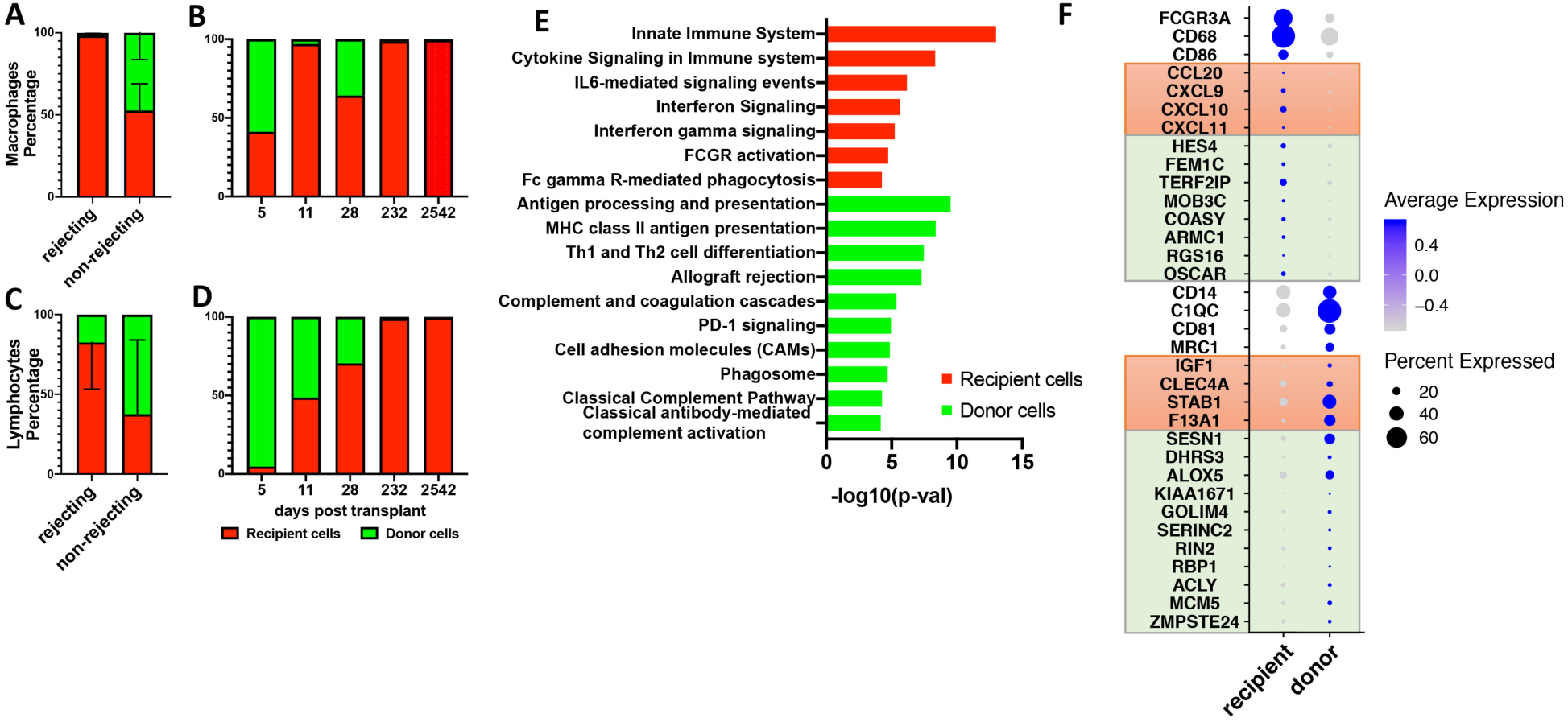
Donor origin Macrophage and Lymphocyte population variations with time and rejection status. A) Macrophages proportions vary according to rejection status. Recipient cell error bars in upwards directions and donor cell error bars downwards direction. B) The proportion of donor to recipient macrophages in each of the 5 biopsies according to time (days) post-transplant. C) The proportions donor and recipient lymphocytes according to rejection status. D) The proportion of donor to recipient T cells in each of the 5 biopsies according to time (days) post-transplant. E) Macrophage pathway analysis by cell origin. Differential gene testing between donor and recipient macrophages reveal genes involved in immune related pathways (ToppGene Suite pathway analysis, Bonferroni p <0.05). F) Dotplot of genes that define donor and recipient macrophages. Orange boxes include genes differentially expressed in ‘classically-activated’ and ‘wound-healing’ macrophages, respectively. Light green boxes include highly expressed genes from spectrum of macrophage phenotypes as described by Xue et al.

Pathway analysis of differentially expressed genes in recipient vs. donor macrophages suggest distinct roles in the alloimmune reaction (Fig. 3E). Il-6 signaling, interferon signaling and FCγR pathway genes are differentially expressed in recipient macrophages. By contrast, complement pathway, antigen presenting and immunoregulatory genes are differentially expressed in donor macrophages. Recipient and donor macrophages do not fall clearly into either M1 or M2 phenotypes (Fig. S4). Recipient macrophages differentially express CD16(*FCGR3A*) and *CD68*, whereas donor macrophages differentially express *CD14* (Fig. 3F). Donor macrophages express genes associated with a wound healing phenotype and recipient macrophages express genes associated with a classically activated macrophage phenotype (Fig. 3F – orange boxes).(17) A transcriptomics based network analysis of human macrophages by Xue *et al* described 10 stimulus specific clusters of activated macrophages that lie along a continuum of M1 to M2 phenotypes.(18) These clusters are represented in both the donor and recipient macrophages (Fig. 3F – green boxes). Using this model top genes expressed by IFNγ and TNF stimulated macrophages, *FEM1C* and *TERF2IP*, are differentially expressed in recipient macrophages. The top gene expressed by IL4 stimulated macrophages, *KIAA1671*, was differentially expressed by donor macrophages.

### Lymphocyte subsets and donor recipient chimerism in the kidney allograft

B cells (n=190), plasma cells (n=231), T cells (n=3067) and mast cells (n=111) could be detected within the lymphocyte cluster (Fig. 4A). Cell types were defined by expression of typical markers including *CD3E* for T cells, *MS4A1* (CD20) for B cells, *SDC1* (CD138) for plasma cells and *TPSAB1* (tryptase) for mast cells (Fig. 4B). Both donor and recipient cells are represented in all lymphocyte subclusters and in mast cells (Fig. 4C) as are rejecting and non-rejecting samples (Fig. 4D).

**Figure 4.**
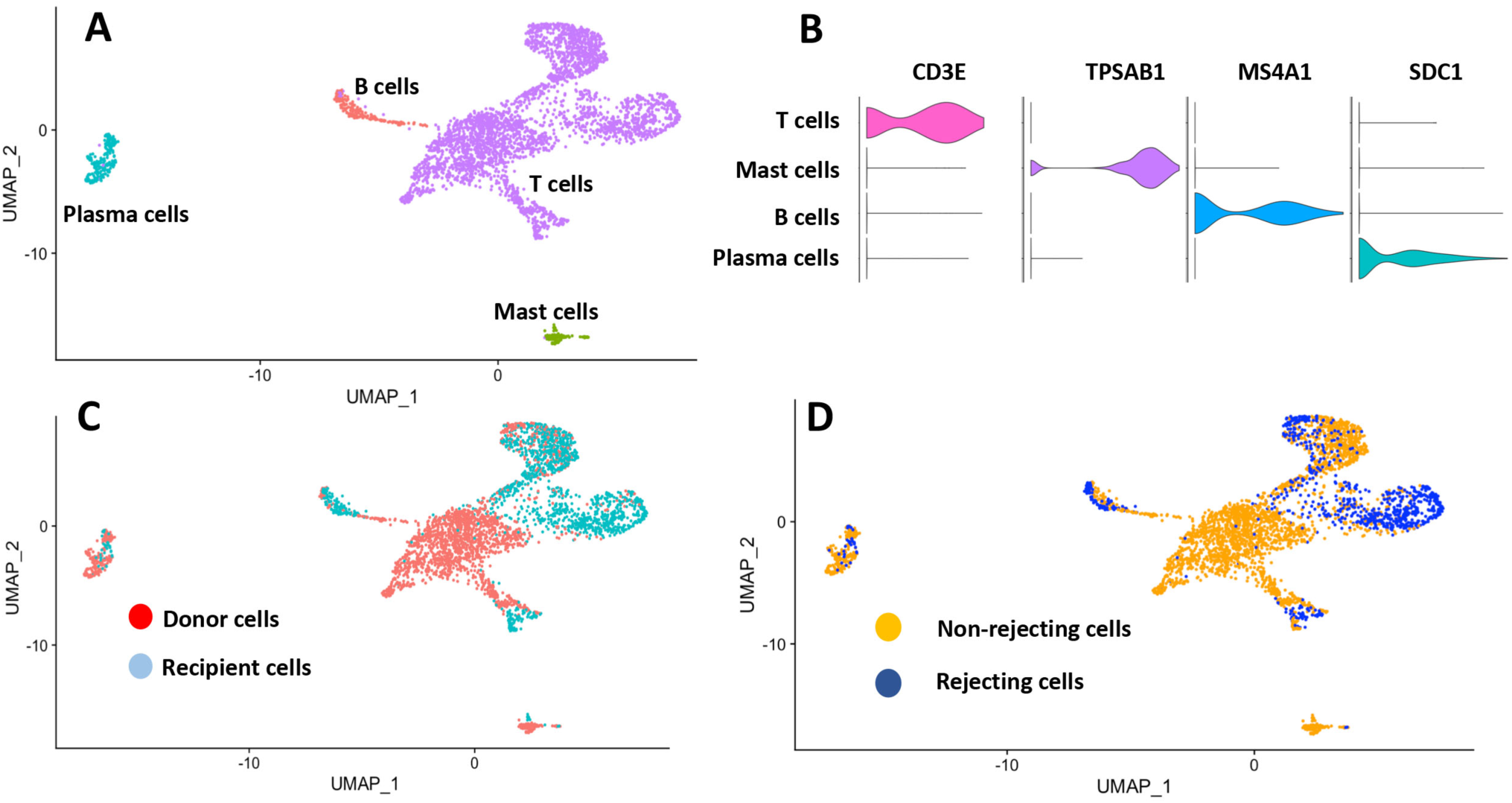
Lymphocytes Subcluster into T Cells, B Cells, Plasma Cells and Mast Cells. A) UMAP visualization of Intragraft lymphocytes subclustered into T cells, B cells, plasma cells and mast cells. B) Violin plot of cell type defining markers, *CD3E* for T cells, *TPSAB1* (tryptase) for mast cells, *MS4A1* (CD20) for B cells and *SDC1* (CD138) for plasma cells. C) UMAP visualization of lymphocytes according to donor or recipient origin and D) rejecting or non-rejecting biopsies.

A differential gene expression list was calculated for recipient T cells versus donor T cells and also for T cells from rejecting biopsies versus T cells from non-rejecting biopsies. Recipient origin T cells correlated highly with rejecting T cell transcripts (Pearson correlation coefficient 0.82). By contrast, donor origin T cells correlated highly with non-rejecting T cell transcripts (Pearson correlation coefficient 0.82) (Fig. 5A). Conversely, T cells from rejecting samples did not correlate with donor origin T cells (Pearson correlation coefficient -0.39) and T cells from non-rejecting samples did not correlate with recipient origin T cells (Pearson correlation coefficient -0.21). Many of the top genes defining recipient T cells and rejecting T cells encode for cytotoxic proteins such as Natural Killer Cell Granule Protein 7 (*NKG7*), granulysin (*GNLY*), granzyme B (*GZMB*), and perforin (*PRF1*) (Fig. 5B). Genes differentially expressed in donor origin T cells and non-rejecting T cells included *IL32, CD52, MIF, S100A6, S100A11, and LTB*. The products of these genes are both pro-inflammatory and anti-inflammatory.

**Figure 5.**
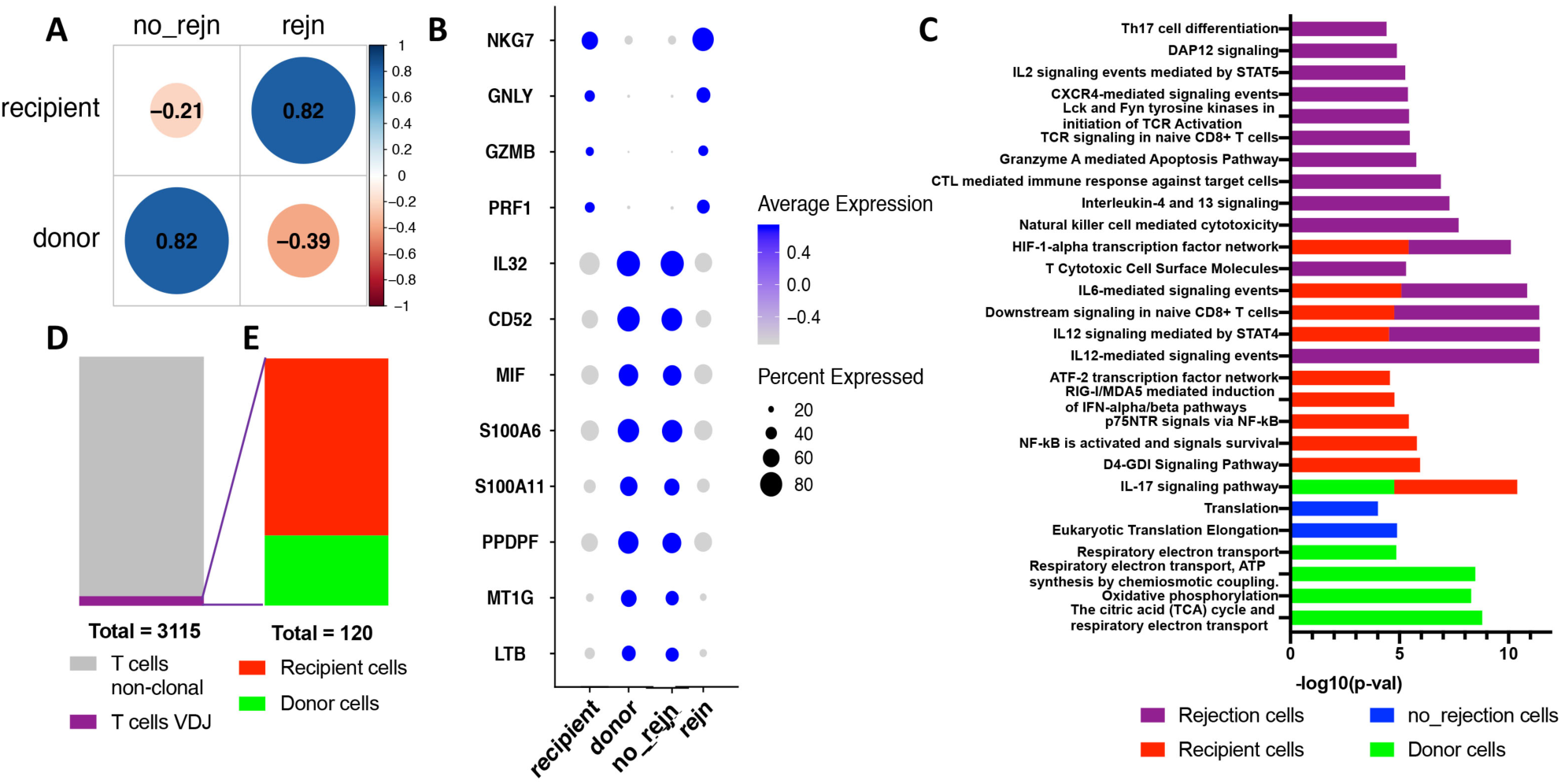
Donor and Recipient Origin T Lymphocyte Relationships with Rejection. A) Pearson correlation analysis of T cells from rejection versus non-rejection samples and recipient versus donor T cells (correlation coefficient are black numbers within circles). B) Select genes from top 20 differentially expressed genes from each group represented in (A), recipient T cells, donor T cells, rejecting T cells and non-rejecting T cells. C) Pathway analysis of significantly differentially expressed genes (Bonferroni p < 0.05).

Pathway analysis using differentially expressed genes defining rejecting, non-rejecting, recipient and donor T cells revealed pro-inflammatory pathways in rejecting and recipient origin T cells (Bonferroni p < 0.05) (Fig. 5C). Pathways common to rejecting and recipient cells were IL-12 signaling through STAT4, IL-6 signaling and downstream signaling in native CD8+ T cells. The HIFα transcription factor pathway was also common to rejecting and recipient T cells. Pathways specific to donor T cells were related to cellular metabolism, specifically oxidative phosphorylation and ATP generation.

### T lymphocyte clonal analysis by VDJ sequencing

We performed single cell sequencing of VDJ rearrangements in T cells from three of the five biopsies in the current dataset (2 rejecting and 1 non-rejecting). Only 120 T cells with VDJ sequence were identified in the final post filter dataset (3.8% of all T cells) (Fig. 5D). There were no T cell VDJ sequences from the non-rejecting biopsy identified in the final dataset. Of the T cells with VDJ rearrangements sequenced 34 cells were donor origin and 86 were recipient origin. All the donor origin VDJ T cells were from one early post-transplantation rejection sample (11 days). The recipient origin VDJ T cells were from a later rejection sample (6 months and 2 weeks) (Fig. 5E).

## Discussion

In this study we used germline DNA sequencing of donor-recipient pairs and scRNA-seq to determine the degree and nature of leukocyte chimerism in the kidney allograft. We studied biopsy samples from three patients with ABMR and two patients with non-rejection causes of AKI. We demonstrate how this approach can accurately assign donor-recipient origin to each cell from a complex tissue such as the kidney and, coupled with transcriptome analysis, can examine the transcriptional differences between cells of different origin.

Although it is known that donor macrophages can be found in the transplanted organ in the early post-transplant period it has been assumed that these cells were transient passengers whose presence declines over time.(19, 20) Furthermore, studies report recipient derived macrophages accumulate during ischemia reperfusion and that following resolution of this injury the macrophage population returned to and remained at low numbers.(21, 22) The current study showed donor derived macrophages can remain within the non-rejecting kidney for up to 28 days despite the infiltration of macrophages associated with the ischemia reperfusion injury that occurs after organ transplantation (Fig. 3A+B). Our data suggests that the donor derived macrophage population may not simply vary as a function of time post transplantation but persist depending on rejection events. Donor macrophage proportions varied according to rejection status rather than time (Fig. 3B). We show that donor derived macrophages can recover post rejection years after transplantation demonstrating the proliferative capacity of these tissue resident donor derived cells (Fig. S3).

Tissue resident macrophages are derived from functionally distinct subsets of cells derived from embryonic and hematopoietic lineages.(23) This distinction suggests that there should be phenotypic heterogeneity of tissue resident macrophages. However, this heterogeneity relies on limited cell markers and animal studies.(14) Such tools to determine macrophage origin in human studies have been limited or not possible. The scRNA-seq approach used in the current study can accurately determine cell origin and has the potential to better define tissue resident macrophage phenotypes. We found functional heterogeneity amongst macrophages in our samples. Functional roles of donor macrophages include wound healing, antigen presentation and negative regulation of T cells through PD-1. Although, classically proinflammatory cytokines are expressed in recipient macrophages, donor derived macrophages appear to include pro-inflammatory processes such as complement activation and coagulation (Fig. 3E+F).

In contrast to macrophages, donor lymphocyte numbers decreased as a function of time and by day 232 there are very few donor origin lymphocytes suggesting that donor to recipient lymphocyte ratios are determined by time post-transplantation (Fig. 3C+D). Recent studies in mice have revealed that memory T cells can reside in non-lymphoid tissue to provide direct antigen recognition and early responses to pathogens.(24) These tissue resident memory T cells (Trm cells) have also been described in kidney.(25) However, the role of Trm cells in transplantation is unknown. In organs with a larger burden of tissue resident T cells such as the lung, donor cells were not detectable after 39 days post-transplantation when assessed by serial transbronchial parenchymal biopsies of lung transplant recipients.(26) The current study demonstrates persistence of a significant number of donor T cells at 28 days post kidney transplantation.

To better understand the functional characteristics of tissue resident T cells we performed an in-depth analysis of differentially expressed genes in donor derived T cells. Interestingly, pathways specific to donor origin T cells (when compared to recipient T cells) are related to glucose metabolism and energy production, specifically oxidative phosphorylation (Fig. 5C). It is known that activated T cells preferentially convert glucose to lactate even when oxygen is not a limiting factor whereas quiescent T cells generate most of their energy through oxidative phosphorylation.(27, 28) The by-products of this increased glycolytic flux in activated T cells are used for the production of nucleotides, fatty acids and amino acids.(29) These data suggest that compared to recipient T cells donor T cells are predominantly quiescent consistent with our findings of donor T cells correlating with T cells from non-rejecting samples (Fig. 5A). Recipient origin T cell state reflects an effector phenotype typical of acute rejection (Fig. 5B+C).

The lymphocyte cluster (Fig. 1) also contained B cells and plasma cells in addition to T cells when subclustering was performed (Fig 4A+B). There was also a small population of mast cells identified. This suggests that mast cells are transcriptionally more similar to lymphocytes in the kidney allograft despite having a myeloid lineage. However, TPSAB1+ mast cells are adjacent to the macrophage cluster in transcriptional space (Fig. S5).

A clonal analysis performed by sequencing the VDJ region of T cells revealed donor derived T cell clones. The significance of this is not known at this time however our ability to identify T cells clones within the kidney, their origin and analyze their transcriptomes adds another category of data previously not available to this field. These donor derived clonal T cells are all from an early post-transplant biopsy and may de passenger T cells that regress with time. Whether the VDJ sequences in these T cells code for donor specific TCRs requires further investigation.

In conclusion, we have combined germline DNA sequencing with scRNA-seq of human kidney transplant biopsies in order to measure the persistence and transcriptional profiles of donor and recipient derived leukocytes in rejection and non-rejecting states. While our sample number is small, we could easily identify leukocytes from both recipient and donor, establishing the feasibility of this approach in transplantation generally.

## Materials and Methods

### Biopsy Samples

Research core biopsy samples were obtained at the time of indication kidney transplant biopsy at Washington University under an Institutional Review Board approved protocol. Biopsy tissue was freshly prepared for study.

### Single Cell isolation and Library Preparation

The renal biopsy was minced into small pieces with a razor blade and incubated at 37°C in freshly prepared dissociation buffer containing 0.25% trypsin and 40 U/ml DNase I. Dissociated cells were harvested every 10 minutes by filtering the cell suspension through a 70-μm cell strainer (pluriSelect) into 10% FBS buffer on ice. The residual biopsy tissue trapped on the cell strainer was dissociated once again with 1 ml dissociation buffer for 10 minutes and passed through the cell strainer into the same FBS buffer from the first collection. We repeated this dissociation procedure three times until most of the tissue had been dissociated into single cells (total dissociation time was approximately 30 minutes). Finally, cells were collected by centrifugation at 400×g for 5 minutes, resuspended in suspension buffer containing 0.1% BSA, and strained through a 40-μm cell mesh (pluriSelect) to further remove cell clumps and large fragments. Cell viability was approximately 90% for the biopsy used in this study as assessed by Trypan Blue staining.

### Bioinformatics and Data Analysis

Cell suspensions were loaded in to 10X Chromium lanes for a target cell capture number of 10,000 cells per lane. Two lanes were used per biopsy. The 10X Chromium libraries were prepared according to the manufacturers protocol. The 5-prime kits were used and T cell VDJ libraries were also prepared according to manufacturer’s protocol.

The Illumina HiSeq platform was used to generate paired end read sequence to a depth of 50,000 reads per cell for gene expression and 5,000 reads per cell for T cell VDJ sequence. The Cellranger v3.0 pipeline was used to generate BAM files and gene expression matrices for each biopsy. Expression matrices were used to generate Seurat objects using Seurat v3 to include cells with genes expressed in at least 3 cells, 200-2500 genes per cell detected and less than 25% mitochondrial genes per cell. Each biopsy was initially clustered using an excessive PC number and high resolution to create a high number of clusters. Clusters containing 2 or more different cell type defining markers were removed as doublet clusters and clusters with a very low average number of genes per cell were removed as debris clusters. An integrated dataset was created using the standard Seurat v3 integration analysis pipeline.

The cor() and corrplot() functions in R were used for Pearson correlation of gene expression between donor and recipient T cells and rejecting and non-rejecting T cells using a significance level of 0.05.

### Donor Recipient Cell Origin – Demuxlet Pipeline(3)

Whole-exome sequencing was performed on each pair of donor and recipient germline DNA from peripheral blood samples associated with each biopsy. Sequencing of each sample was performed targeting a coverage depth of 50X. Alignment was performed using BWA-MEM. MarkDuplicates with Picard followed by BaseQualityRecalibrate (BQSR) with GATK was then performed in accordance with the functional equivalence paper published by the Centers for Common Disease Genomics (CCDG).(30) Files were then converted to CRAM format. Variant call format files (.vcf) were created using GATK best practices filtering for high quality common variants (allele frequency > 0.05). Each combined donor and recipient .vcf and aggregated .BAM file from Cellranger output was used as input for the demuxlet pipeline using demuxlet default parameters and alpha of 0, 0.5 and 1. Each cell in the final integrated data set was annotated as donor or recipient using the best column from the demuxlet output data.

### Pathway Analysis

Significant differentially expressed genes (adjusted p value < 0.05) from the Seurat findmarkers() (minimum percentage of cells with gene = 0.25, log fold change threshold of 0.25) were analyzed using the Toppfun online tool. Pathways with a Bonferroni p value above 0.05 were removed from analysis. For T cell pathway analysis, pathways for which only CD3 and CD2 gene expression was common to the reference pathway gene set were excluded from analysis.

### T Cells VDJ Immune Cell Profiling

T cell VDJ sequence was analyzed using VDJ Loupe Browser. Clones were defined as cells with complete VDJ sequence from both alpha and beta chains. Using cell barcodes T cell clones were identified in the final integrated dataset.

## Acknowledgments

This work was supported by the McDonnell Genome Institute 2018 Symposium Pilot Project Fund at Washington University as well as the NIH/National Center for Advancing Translational Sciences (NCATS) grant UL1TR002345 (to C.F., R.F., A.F.M. and B.D.H) and. NIH/NIDDK grants 1K08DK120953 (to A.F.M.) and DK103740 and DK107374 (to B.D.H).

