## Supplemental Material for "Harnessing Expressed Single Nucleotide Variation and Single Cell RNA Sequencing to Define Immune Cell Chimerism in the Rejecting Kidney Transplant"

Benjamin D. Humphreys, MD, PhD

Division of Nephrology

Washington University School of Medicine

660 S. Euclid Ave., CB 8129

St Louis, MO 63110

**This PDF file includes:**

Supplementary text
Figures S1 to S4
Table S1

### Supplementary Information Text

### Methods

#### Biopsy Samples

Research core biopsy samples were obtained at the time of indication kidney transplant biopsy at Washington University under an Institutional Review Board approved protocol. Biopsy tissue was placed in Wisconsin buffer and placed on ice prior to immediate preparation for study.

#### Single Cell isolation and Library Preparation

The renal biopsy was minced into small pieces with a razor blade and incubated at 37°C in freshly prepared dissociation buffer containing 0.25% trypsin and 40 U/ml DNase I in phosphate buffered saline. Dissociated cells were harvested every 10 minutes by filtering the cell suspension through a 70-µm cell strainer (pluriSelect) into 10% FBS on ice. The residual biopsy tissue trapped on the cell strainer was dissociated once again with 1 ml dissociation buffer for 10 minutes and passed through the cell strainer into the same FBS buffer from the first collection. We repeated this dissociation procedure three times until most of the tissue had been dissociated into single cells (total dissociation time was 30-40 minutes). Finally, cells were collected by centrifugation at 400×g for 5 minutes, resuspended in suspension buffer containing 0.1% BSA, and strained through a 40-µm cell mesh (pluriSelect) to further remove cell clumps and large fragments. Cell viability was approximately 90% for the biopsy used in this study as assessed by Trypan Blue staining.

**Generation of cDNA libraries for scRNA-seq**
Cell suspensions were loaded in to 10X Chromium lanes for a target cell capture number of 10,000 cells per lane. Two lanes were used per biopsy. The 10X Chromium libraries were prepared according to the manufacturers protocol. The 5-prime kits were used and T cell VDJ libraries were also prepared according to manufacturer's protocol.

**Next Generation Sequencing of scRNA-seq libraries**
The concentration of each prepared 10x single cell library was accurately determined through qPCR utilizing the KAPA library Quantification Kit according to the manufacturer's protocol (Roche) to produce cluster counts appropriate for the Illumina NovaSeq6000 instrument. Paired end sequence data (2x150) on the S4 flow cell was generated targeting 50,000 total read pairs per cell for gene expression and 5,000 total read pairs per cell for VDJ sequence. The Cellranger 3.0 pipeline was used to generate BAM files and gene expression matrices for each biopsy.

**IDT Exome Sequencing (Whole-exome Sequencing) Methods**
250ng of genomic DNA was fragmented on the Covaris LE220 instrument targeting 250bp inserts. Automated dual indexed libraries were constructed with the KAPA HTP library prep kit (Roche) on the SciClone NGS platform (Perkin Elmer). Ten libraries were pooled at an equimolar ratio by mass prior to the hybrid capture targeting a 5µg library pool. The library pool was hybridized with the xGen Exome Research Panel v1.0 reagent (IDT Technologies) that spans 39 Mb target region (19,396 genes) of the human genome. The libraries were hybridized for 16-18 hours at 65°C followed by stringent washing to remove spuriously hybridized library fragments. Enriched library fragments were eluted with streptavidin-coated magnetic beads and amplified with KAPA HiFi Polymerase prior to sequencing. PCR cycle optimization was performed to prevent over amplification of the libraries. The concentration of each captured library pool was accurately determined through qPCR utilizing the KAPA library Quantification Kit according to the manufacturer's protocol (Roche) to produce cluster counts appropriate for the Illumina NovaSeq6000 instrument. Approximately 5Gb of paired end sequence data (2x150) on the S4 flow cell was generated targeting 50x coverage per sample. Alignment was performed using BWA-MEM. MarkDuplicates with Picard followed by BaseQualityRecalibrate (BQSR) with GATK was then performed in accordance with the functional equivalence paper published by the Centers for Common Disease Genomics (CCDG)(1). Files were then converted to CRAM format.

**Generation of Seurat Objects**
Cellranger output files were used to generate count matrices using Read10X() in R. Seurat v3 CreateSeuratObject() was then used and the final object was refined to include cells with genes expressed in at least 3 cells, 200-2500 genes per cell detected and less than 25% mitochondrial genes per cell. The resolution and number of principle components used in the final Seurat object for each biopsy varied. Each biopsy was pre-clustered using an excessive PC number and high resolution to create a high number of clusters. Cluster defining genes based on the FindAllMarkers() function were then examined. Clusters containing 2 or more different cell type defining markers were removed as doublet cells and clusters with a very low average number of genes per cell were removed as debris clusters if no single cluster defining gene was identified. Once doublet clusters were removed the final object was recreated using a lower resolution and PC number.

**Generation of Integrated Seurat Object**
An integrated dataset was created using the standard Seurat v3 integration analysis pipeline. Each input object was log-normalized prior to variable feature selection based on a variance stabilizing transformation to find the 2000 most variable genes. 30 principle components were calculated for PC analysis and the final integrated object was constructed using 11 PCs.

**Donor Recipient Cell Origin – Demuxlet Pipeline(2)**
Variant call format files (.vcf) were created from the CRAM output file from the WES pipeline using GATK best practices. A single .vcf file was created for each donor-recipient pair. These files were filtered for high quality common variants (allele frequency > 0.05). An aggregated .BAM file was created from the .BAM files from each of the 2 single cell 10X lanes used for each biopsy. The aggregated .BAM file and the .vcf file from each biopsy was used as input for the demuxlet pipeline using demuxlet default parameters and alpha of 0, 0.5 and 1. Each cell in the final integrated data set was then annotated as donor or recipient using the best column from the demuxlet output data.

**Identification of differentially expressed genes**
Differentially expressed genes were identified by comparing the transcriptional profile of donor and recipient cells within the macrophage or T cell clusters in the integrated dataset using FindMarkers() with logfc.threshold=0.25 and min.pct=0.25. Also, differentially expressed genes were identified by comparing the transcriptional profile of T cells from rejecting biopsies and T cells from non-rejecting biopsies using the same method.

**Correlation plots and DotPlots**
The cor() and corrplot() functions in R were used for a Pearson correlation of gene expression between donor and recipient T cells and rejecting and non-rejecting T cells using a significance level of 0.05. The Seurat function DotPlot with default parameters was used to compare gene expression of top genes between donor and recipient cells and rejecting and non-rejecting cells.

**Pathway Analysis**
Significant differentially expressed genes (adjusted p value < 0.05) from the Seurat function FindMarkers() (minimum percentage of cells with gene = 0.25, log fold change threshold of 0.25) were used for pathway analysis. GO analysis was performed on the differentially expressed genes using the ToppGene Suite (https://toppgene.cchmc.org). Non-significant GO terms (Bonferroni p value above 0.05) were removed from analysis. For T cell pathway analysis, pathways for which only CD3 and CD2 gene expression was common to the reference pathway gene set were excluded from analysis. Significant enriched GO terms (defined by defined by Bonferroni P-value <0.05) were summarized by REVIGO(3) and visualized by treemap R package.

**T Cells VDJ Immune Cell Profiling**
T cell VDJ sequence was analyzed using VDJ Loupe Browser. Clones were defined as cells with complete VDJ sequence from both alpha and beta chains. Using cell barcodes T cell clones were identified in the final integrated dataset.

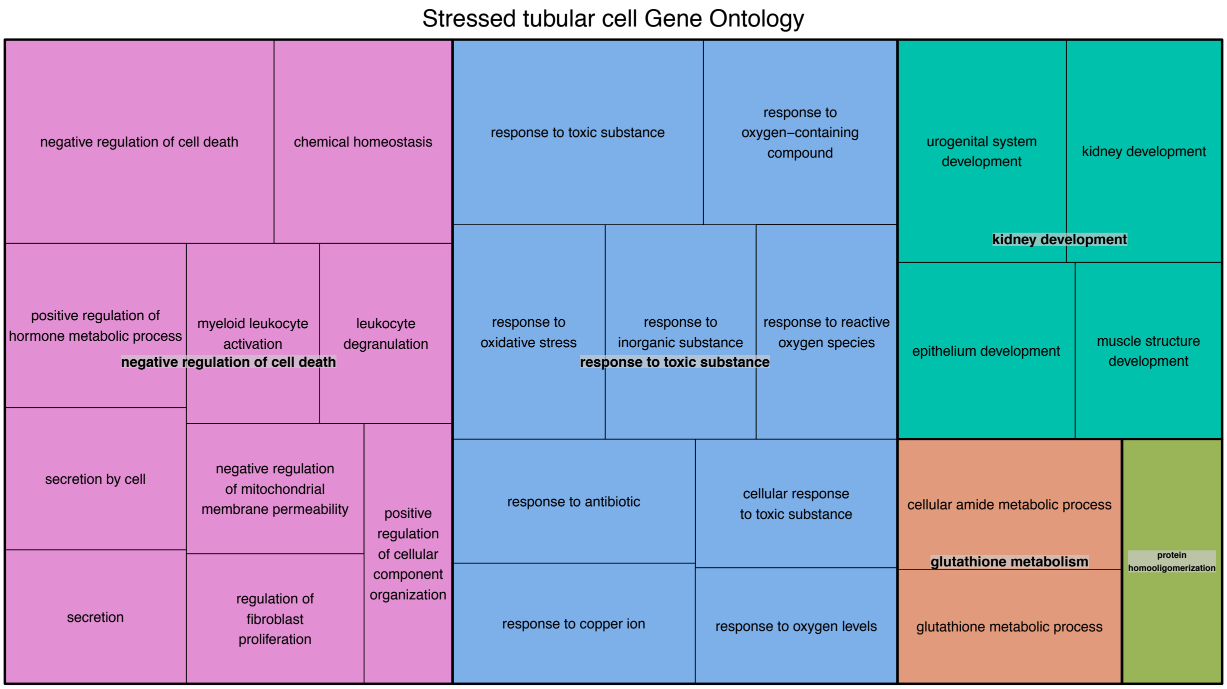


Figure S1. Gene ontology terms for differentially expressed genes that define stressed tubular cell cluster. Top terms were negative regulation of cell death and responses to external toxins including oxidative stress.

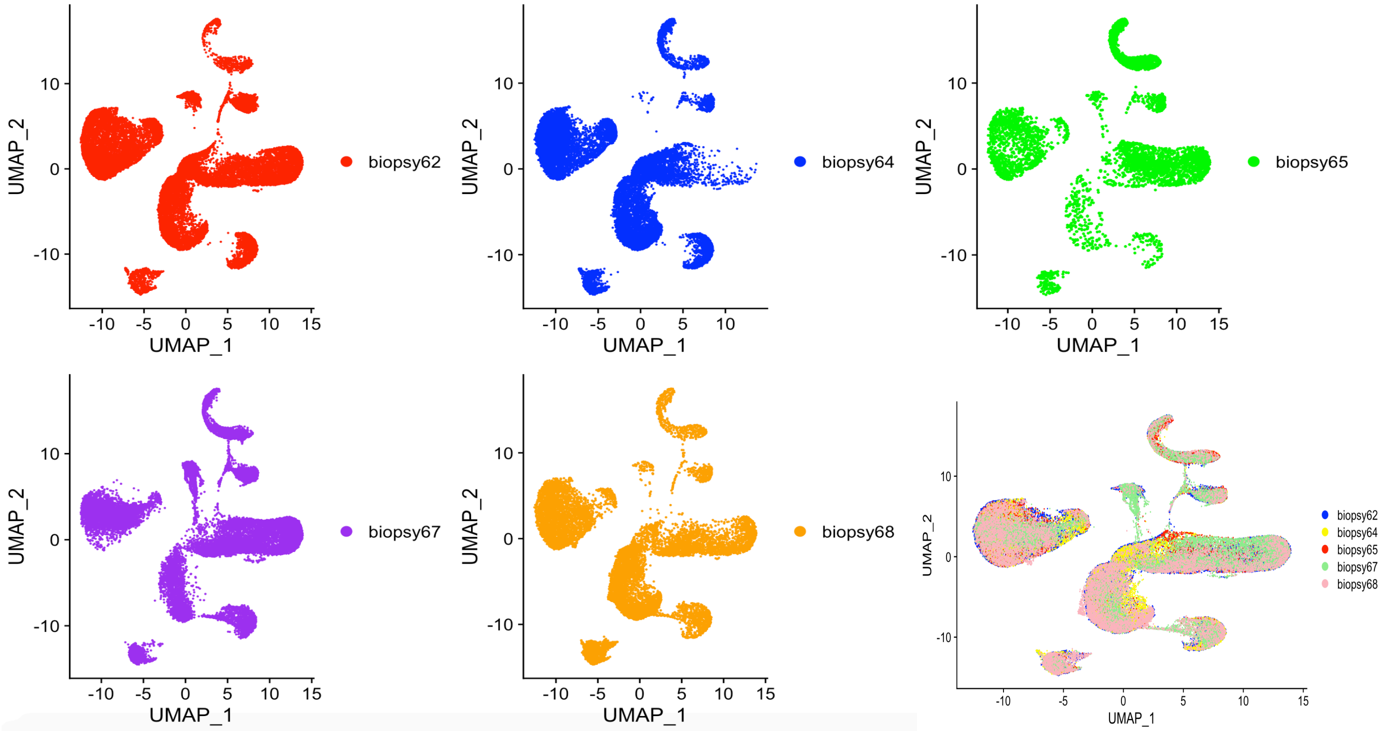


Figure S2. A+B+C+D+E) UMAP visualization of each individual biopsy in the dataset demonstrating equal coverage of all cell clusters. F) UMAP visualization of the whole dataset grouped by original biopsy identity.

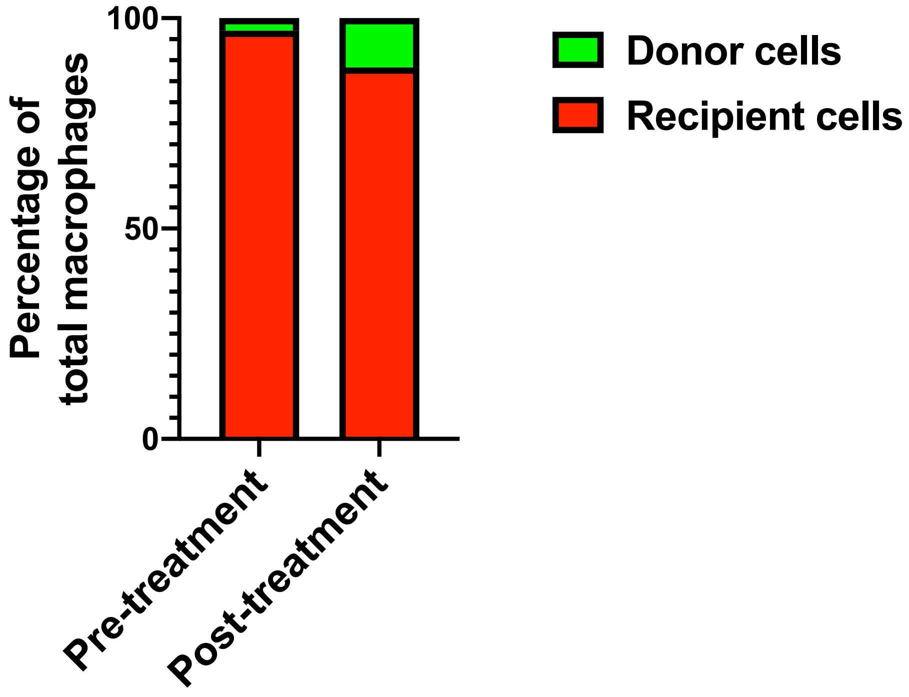


Figure S3. Serial biopsies in the same patient at rejection (ABMR) diagnosis (Pre-treatment) and one month post treatment demonstrates donor macrophage proportion increases significantly post rejection treatment. Pre-treatment biopsy was performed 2years 7 months (945 days) post-transplant and the post treatment biopsy was performed one month after the first biopsy (974 days).

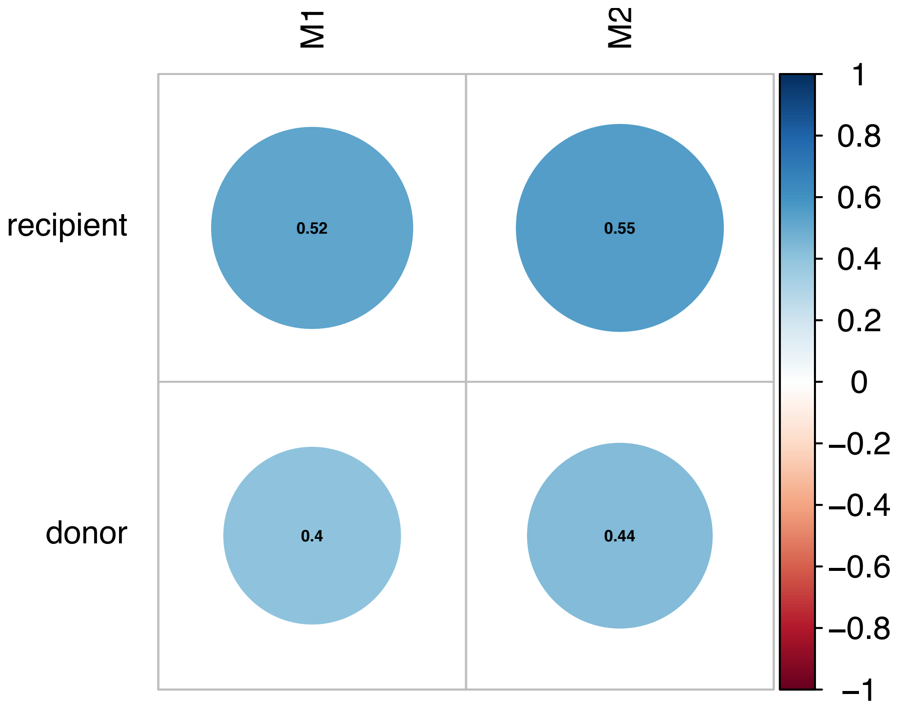


Figure S4. gene expression from donor and recipient macrophages were correlated with human M1 (n=4) and M2 (n=4) macrophage bulk RNA-seq gene expression external datasets. K. Y. Gerrick *et al.*, Transcriptional profiling identifies novel regulators of macrophage polarization. *PloS one* **13**, e0208602 (2018).

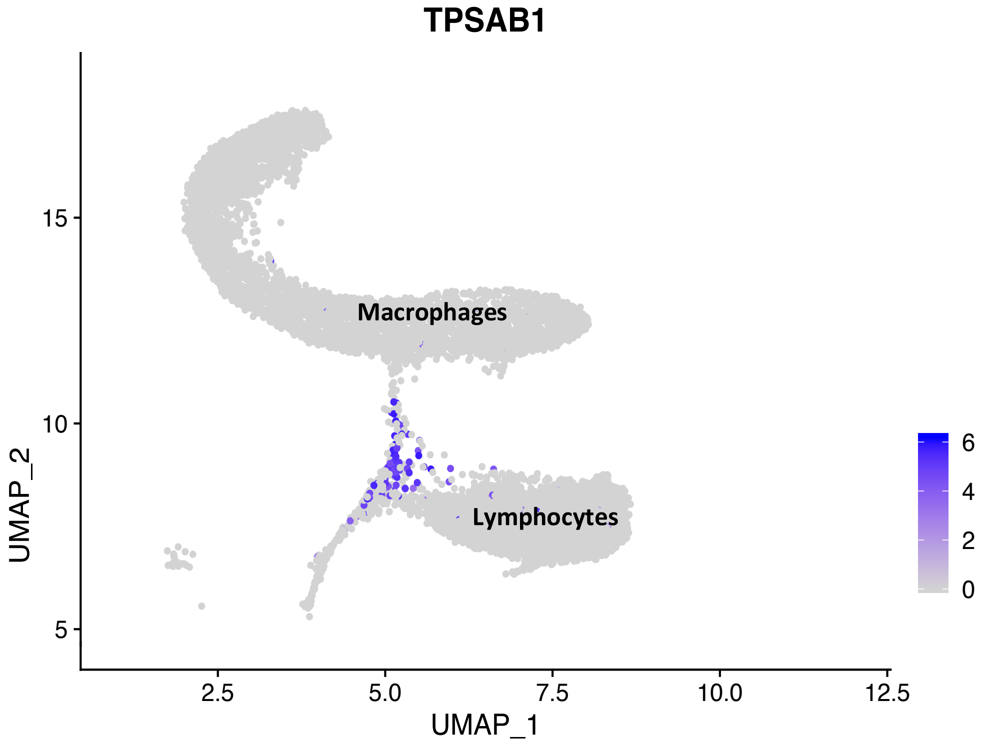


Figure S5. UMAP visualization of macrophages and lymphocytes relative to TPSAB1 (tryptase) positive Mast cells (purple)

### Table S1. Clinical Characteristics of the 5 Biopsies

| Rejection Status | Patient Age (yrs) | Time post Transplant (days) | Biopsy Indication | Biopsy diagnosis | Donor Specific Antibodies |
|---|---|---|---|---|---|
| No Rejection | 53 | 5 | AKI | Acute tubular injury, No rejection | No<br>(DR1 1500mfi) |
| Rejection | 38 | 11 | AKI and proteinuria 2.1g | ABOi rejection, Thrombotic microangiopathy . No cellular rejection | Yes<br>anti-A 1:32 |
| No Rejection | 55 | 28 | AKI and proteinuria 3.1g | Acute tubular injury. No rejection | No |
| Rejection | 75 | 232 | AKI and previous ABMR | Active ABMR, C4d is diffusely positive | Yes<br>B8(1190mfi);<br>DQA1*05:01(17846mfi);<br>DQA1*03(5153mfi) |
| Rejection | 35 | 2542 | AKI and proteinuria 6.7g | Chronic Active ABMR, C4d is focally positive. IgA nephropathy consistent with recurrence | Yes<br>DR1(5834mfi);<br>DR53(25460mfi);<br>DQ2(7623) |
